# High contrast staining for serial block face scanning electron microscopy without uranyl acetate

**DOI:** 10.1101/207472

**Authors:** Adolfo Odriozola, Jaime Llodrá, Julika Radecke, Céline Ruegsegger, Stefan Tschanz, Smita Saxena, Rohr Stephan, Benoît Zuber

## Abstract

Serial block face scanning electron microscopy (SBFSEM) is an increasingly popular method for investigating the three-dimensional ultrastructure of large biological samples. Prior to imaging, samples are typically chemically fixed, stained with osmium and uranyl acetate, and subsequently embedded in resin. The purpose of staining is to provide image contrast and reduce specimen charging under the electron beam, which is detrimental to the quality of imaging. Obtaining, using, and disposing of uranyl acetate is getting increasingly cumbersome in many countries due to new regulations on the handling of radioactive substances. Therefore, we developed an alternative staining procedure that does not rely on the use of uranium or any other radioactive substance. This procedure provides excellent contrast and efficiently reduces specimen charging.

## 2 Introduction

By providing access to cell and tissue ultrastructure at nanometer resolution, transmission electron microscopy (TEM) has revolutionized cell biology in the mid-20th century. Yet classical TEM only provides 2-dimensional images, which hinders the assessment of the 3-dimensional cellular structure of the tissue under investigation. TEM analysis of serial sections has alleviated this problem but the throughput for producing, imaging, and processing serial sections of useful sample volume is very low. Recently two scanning electron microscopy (SEM) modalities have been introduced where a resin-embedded specimen block is placed in the microscope and its surface is repeatedly and automatically 'shaved’ and imaged, producing serial images at a much higher throughput [1]. ‘Shaving’ is achieved by using either a focused ion beam (FIBSEM: focused ion beam SEM) or an ultramicrotome with a diamond knife, which is mounted inside the microscope specimen chamber (SBFSEM: serial block face SEM). SBFSEM can image larger volumes than FIBSEM although at somewhat lower resolution (20 to 30 nm vs. 5 to 10 nm) [1, 2]. Note that these numbers refer to effective resolution and not to pixel size, which is often mistaken for resolution in the FIBSEM and SBFSEM fields. Resin-embedded specimens must be contrasted by heavy metal staining. While for TEM imaging specimen preparation typically involves a combination of in-block and on-section staining, FIBSEM and SBFSEM imaging depends exclusively on in block staining, which renders the procedure more difficult, since in FIBSEM and SBFSEM the block face, and not sections, is imaged. Aside its function in contrast establishment, heavy metal staining renders specimens electrically conductive. This counteracts specimen charging occurring under the scanning electron beam thereby leading to image distortion. Charging is a particular concern in SBFSEM, where it is inherently more prone to occur than in FIBSEM. Indeed it is thought that ion beam scanning neutralizes the charges induced by the electron beam scan. Furthermore charging has another deleterious consequence in SBFSEM as it weakens the resin and leads to fragmentation of the block surface during sectioning. A highly efficient heavy metal staining procedure is therefore of critical importance for samples to be imaged by SBFSEM.

Most staining protocols for SBFSEM consist of successive incubations with reduced osmium, a mordant for osmium, osmium tetroxide, uranyl acetate, and lead aspartate [3, 4, 5, 6]. It is generally considered that osmium mainly stains membranes, whereas uranyl stains proteins and chromatin and lead enhances uranyl staining [7]. Working with uranyl acetate has become increasingly cumbersome in recent years due to the introduction of stricter regulations on radioactive material in several countries. In addition, some research institutions enforce more rigorous rules than the legal requirements in term of radioactive waste disposal. This results in time and cost intensive procedures. We therefore sought to develop an alternative staining protocol for SBFSEM that does not rely on the use of radioactive material. Lanthanides have been used as alternative on-section stain for TEM and have provided similar staining properties to uranyl acetate [8, 9]. On this basis we developed a protocol that uses lanthanides, namely samarium and gadolinium salts, instead of uranyl acetate during bloc staining. We tested the protocol on various samples. The results show that contrast is at least as good or improved, whereas charging is reduced.

## 3 Results

### 3.1 Cerebellar cortex

We sought to develop a bloc staining protocol free of radioactive uranyl acetate for SBF-SEM (see Methods). Instead of uranyl acetate, we used gadolinium triacetate and samarium triacetate, which were previously shown to be efficient staining agents for ultrathin sections [8]. We tested the protocol on mouse cerebellar cortex. The contrast provided by this procedure was excellent (Figure 1). The two membranes composing the nuclear envelope, as well as the two membranes of endoplasmic reticulum cisternae were clearly resolved (Figure 1B and S1 Movie). On the other hand, nuclear membranes of specimens of samples processed with the ROTO protocol (using uranyl acetate as defined under Methods) appeared as a single line (Figure 1C). Furthermore, with the ROTO protocol, nuclear pores were poorly visible in grazing section and practically unresolved in crosssection, whereas they were obvious with lanthanide staining, both in grazing section and in cross-section (arrows in Figure 1B and C, S1 Movie). Contrast after lanthanide staining was such that the nuclear envelope and nuclear pores could be rendered in 3-dimension without the need of manual segmentation (Figure 1D). From such representation it is obvious that Purkinje cell nuclear pores are frequently found in small clusters of pores, consistent with a previous report [10]. Heavy electron radiation necessary to obtain high resolution SBFSEM images often generates chemical modification of the specimen leading to severe sectioning artefacts, which preclude acquisition of 3-dimensional stacks at high resolution [11]. These damages are especially pronounced in samples with low electrical conductivity. From the lanthanide-based protocol processed samples we could record high resolution stacks of hundreds of images with a slice thickness of 50 nm without any noticeable damages, indicating efficient staining resulting in good conductivity (S1 Movie). Moreover mitochondria cristae as well as synaptic vesicles were well resolved after lanthanide staining (Figure 2A, S1 Movie) whereas they were ill-defined after ROTO staining (Figure 2B).

**Figure 1.**
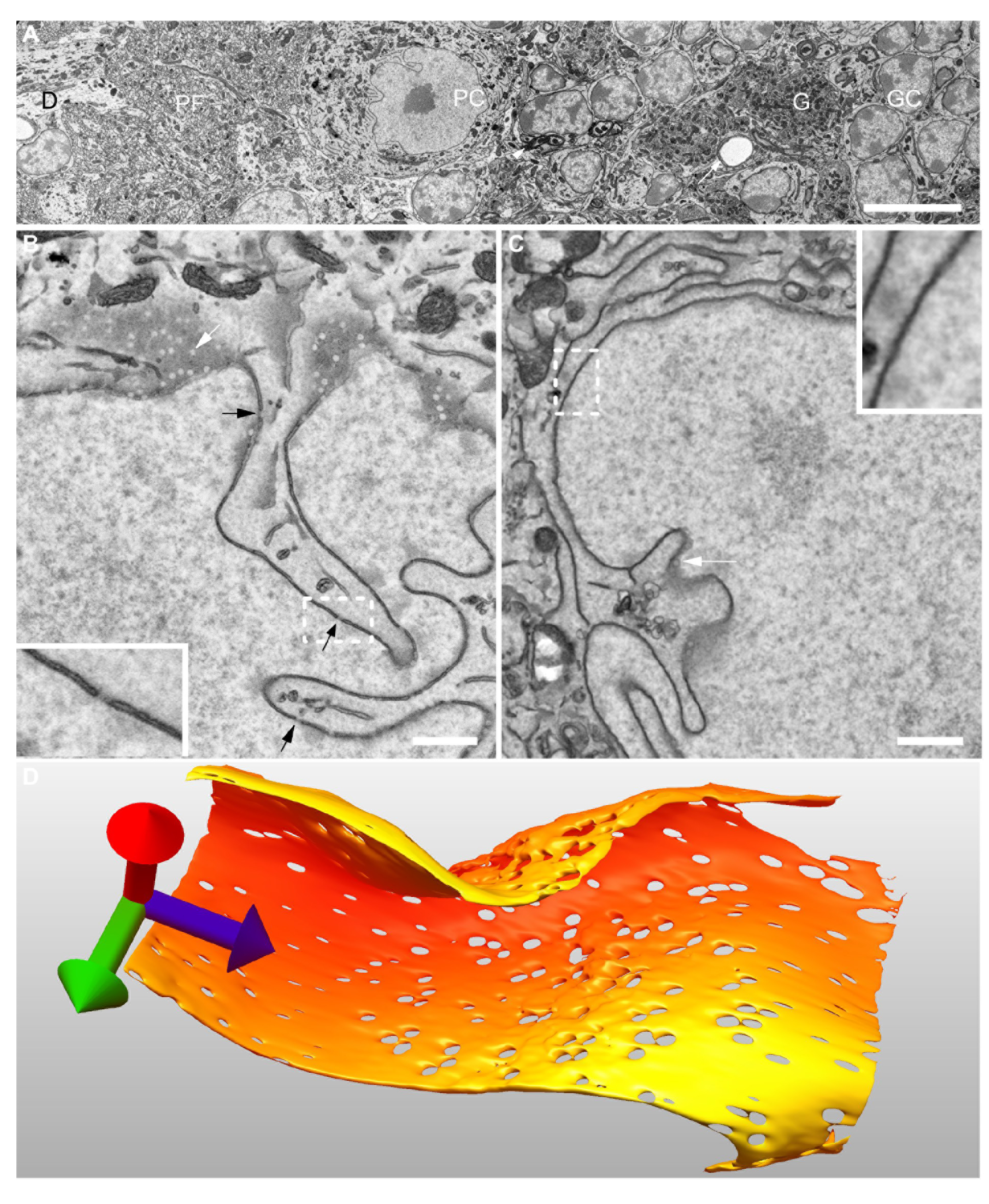
SBFSEM images of cerebellar cortex. A: Overview of cortex prepared with the lanthanide-based protocol shows excellent contrast. (D, Purkinje cell dendrites; PF, parallel fibers, PC, Purkinje cell body; G, cerebellar glomerulus; GC, granule cell body). B: High magnification view of a Purkinje cell nucleus prepared with the lanthanide protocol showing resolved nuclear membranes and nuclear pores in cross-section (black arrows and inset) and in grazing section (white arrows). C: High magnification view of a Purkinje cell nucleus prepared with ROTO. The nuclear envelope appears as a single line (inset). Nuclear pores are only visible in grazing section (white arrow). D: Isosurface rendering of a portion of a Purkinje cell nuclear envelope prepared with the lanthanide protocol. The blue vector is perpendicular to the block face. Original pixel sizes: 40 nm (A), 3 nm (B), 6 nm (C), Bars: 10 μm (A), 1 μm (B, C)

**Figure 2.**
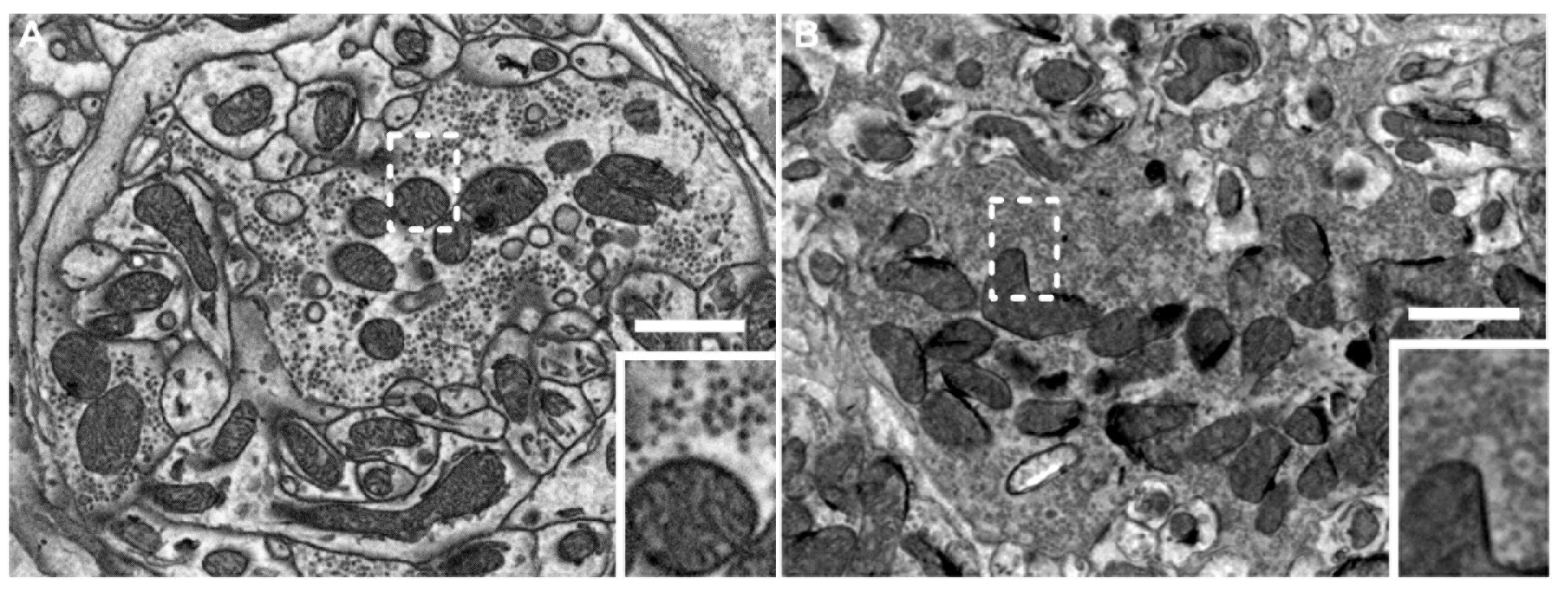
SBFSEM images of cerebellar cortex mossy fiber synapses. A: High magnification view of a mossy fiber synapse in a cerebellar glomerulus prepared with the lanthanide protocol. Synaptic vesicles, as well as mitochondrial cristae are clearly resolved (inset). B: High magnification image of a mossy fiber synapse prepared with the ROTO protocol. Synaptic vesicles are less well resolved and cristae are barely visible. Original pixel size: 5.5 nm. Bars: 1 μm.

**Figure 3.**
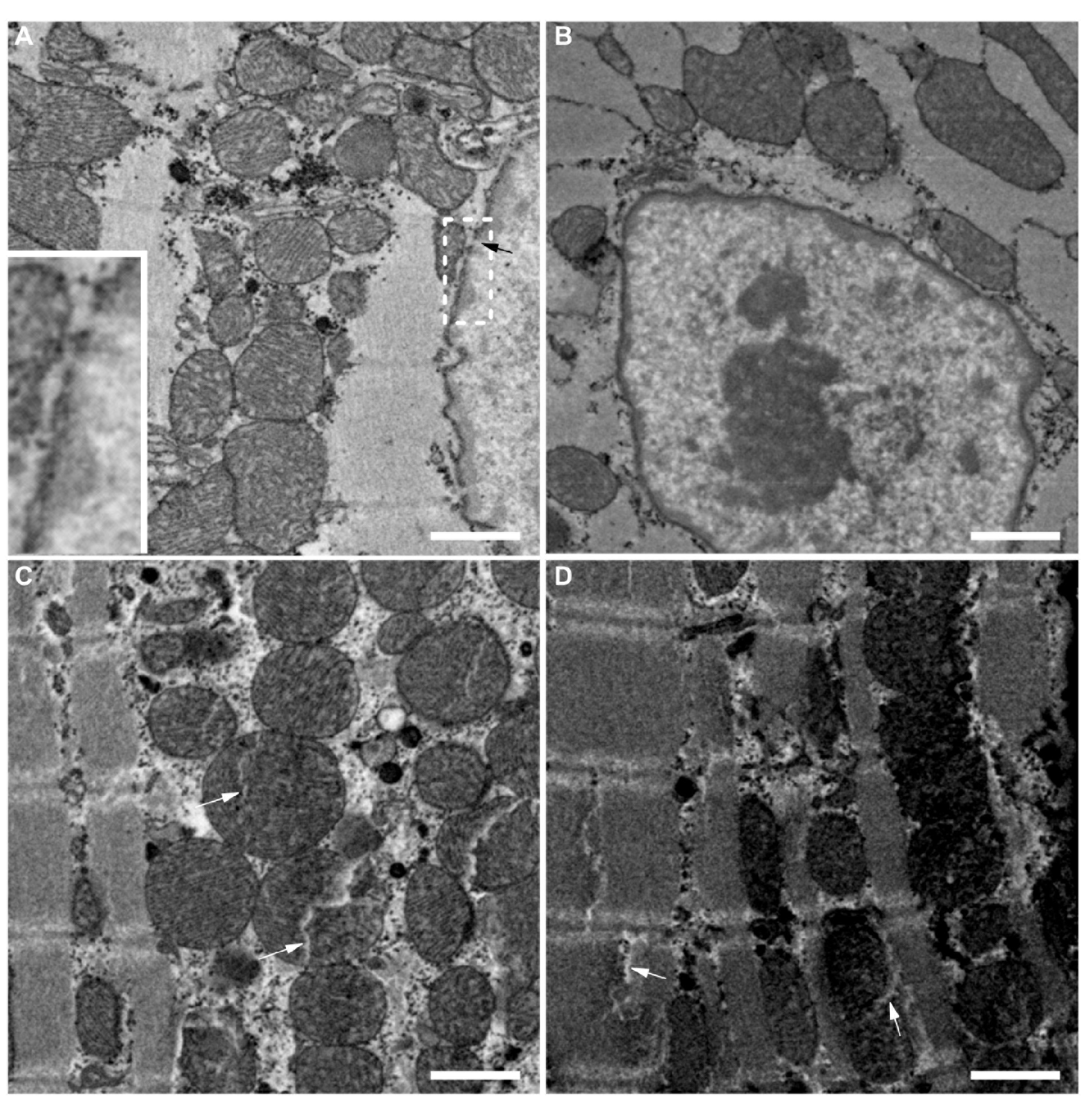
High magnification SBFSEM images of cardiomyocytes. A: Sample prepared with the lanthanide protocol. Cristae as well as nuclear pores (black arrow, inset) were well resolved. B: Sample prepared with the lanthanide protocol where the osmication steps were performed at room temperature. Nuclear pores are not resolved and cristae are hardly visible. C and D: Sample prepared with the lanthanide protocol with longer osmication steps at 50°C. These two images show two fields of view, which are 6 μm apart. The image in C shows good contrast whereas the area in D is too strongly stained. In the latter, mitochondria appear so dark that cristae are not visible. White arrows indicate cracks. Original pixel sizes: 6 nm (A-D). Bars: 1 μm (A-D).

### 3.2 Myocardium

We applied the lanthanide-based protocol to rat myocardium and we decided to assess on this tissue if the duration of the osmication steps performed at high temperature could influence the staining quality. First we applied the exact same osmication steps as for cerebellum (OsO_4_ and potassium ferrocyanide incubation for 45 minutes at room temperature and then for 15 minutes at 50 °C, pyrogallol for 15 minutes at room temperature and for 5 minutes at 50 °C, and OsO_4_ for 22 minutes at room temperature and 8 minutes at 50 °C). As for cerebellum, the mitochondria cristae, nuclear membranes, and nuclear pores were resolved (Figure 3A). Furthermore, myofibrils and the sarcoplasmic reticulum were visible. These images, as those of cerebellar cortex, were acquired under high vacuum (2 mPa). This implies that the samples were electrically conductive, in agreement with their strong staining. When all osmication steps were performed at room temperature (OsO_4_ and potassium ferrocyanide incubation for 60 minutes, pyrogallol for 20 minutes, and OsO_4_ for 30 minutes, all at room temperature), myocardium samples were too lightly stained and not sufficiently conductive (Figure 3B). Hence they had to be imaged under low vacuum conditions (40 Pa) to prevent charging. The individual membranes of the nuclear envelope could not be resolved at all. Nuclear pores were hardly visible and mitochondria cristae were far less well resolved than when osmication was performed at 50 °C. Given the improved quality provided by osmication performed at 50 °C, we decided to increase the duration of the steps performed at 50 °C with the aim of further improving contrast (incubation in OsO_4_ and potassium ferrocyanide for 60 min at 50 °C, incubation with pyrogallol for 20 minutes at 50 °C, and second incubation in OsO4 for 30 min at 50 °C). Some areas of the samples were well contrasted (Figure 3C) whereas adjacent areas were too strongly stained (Figure 3D). Consequently the contrast was low since the material was mostly dark. Note that in these, several cracks were visible indicating a too strong staining [12]. Even in the less strongly stained areas cracks were visible in mitochondria and their cristae were poorly resolved (Figure 3C). Taken together these results demonstrate that the duration of the osmication steps can be optimized in order to obtain best contrasting and conducting performances.

### 3.3 Lung

Finally, we applied our protocol to rat lung and compared it to the ROTO procedure. The alveolar air compartment represents a large proportion of the sample and after specimen preparation it is filled with unstained resin. Consequently large areas of block surface are not conductive and imaging must be performed under low vacuum conditions (60 Pa). The lanthanide protocol provided a better contrast. For example, vesicles and other membranous compartment stood out better over cytoplasmic background and the contrast between hetero-and euchromatin was stronger (Figure 4). Lamellar bodies were saturated with stain after the ROTO procedure whereas they were more lightly stained and hence their inner structure was better resolved after the lanthanide protocol.

**Figure 4.**
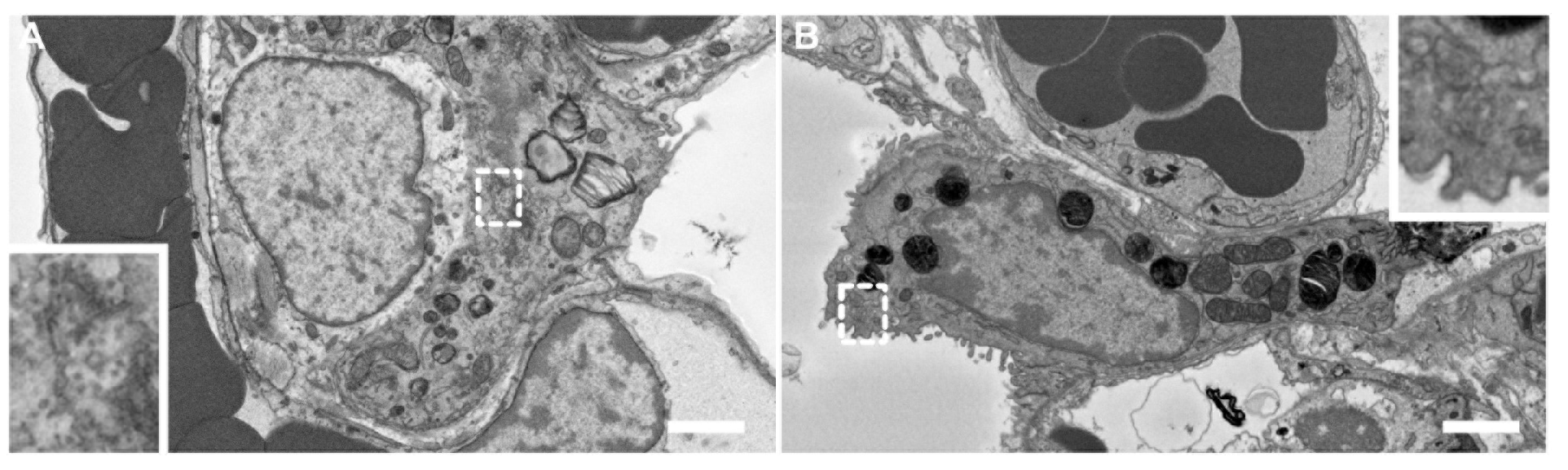
SBFSEM images of lung alveoli processed with the lanthanide protocol (A) or the ROTO protocol (B). On each image, a type II epithelial is visible. The contrast in these cells is slightly better after the lanthanide protocol (see insets). Original pixel sizes: 5.7 nm (A), 6 nm (B). Bars: 2 μm (A-D).

## 4 Discussion

SBFSEM has gained a lot of interest in the last couple of years. It allows fast acquisition of 3-dimensional ultrastructural information. Uranyl acetate is used in most procedures for sample staining prior to SBFSEM. Uranyl acetate is getting more and more difficult to obtain due to increasingly stricter regulations on radioactive substances. Hence we sought for a non-radioactive substitute that would efficiently stain biological samples for SBFSEM. Lanthanides salts have been successfully used to stain ultrathin sections for TEM [8, 9]. As they are not radioactive, we decided to perform en bloc staining with these salts. We applied them after osmication steps. Furthermore, we optimized osmication by performing some steps at 50 °C. This resulted in excellent contrast allowing for example to clearly resolve the two nuclear membranes, nuclear pores, and mitochondrial cristae. In our hands, our new protocol provided significantly better staining than the uranyl-based ROTO protocol. Most samples could be imaged under high vacuum, which is a prerequisite to reach the best possible resolution. Samples that contained a lot of non-stainable volume (e.g., air compartment in lung alveoli) needed to be imaged under low vacuum to prevent charging. Yet contrast in those samples was still very good. The protocol that we present here should be applicable to most biological samples. For each new type of specimen, we recommend to optimize the duration of the steps performed at 50 °C as we demonstrated for myocardium. We anticipate that combining the new lanthanide-based staining protocol and sample embedding in conductive resins [6] will contribute to achieving optimal results.

## 5 Methods

All incubations in fixative were done at 4 °C. All other steps were performed at room temperature except if indicated differently.

### 5.1 Chemicals, cell culture, animals

If not otherwise specified, chemicals were purchased from Sigma-Aldrich (Buchs, Switzerland). OsO_4_ and uranyl acetate were obtained from Electron Microscopy Sciences (Hatfield, PA, USA). Lung and heart samples were obtained from adult Wistar rats. Cerebellum samples were obtained from adult C57BL/6 mice. Milli-Q water was used to prepare all solutions and to perform all the rinses described in the steps below.

### 5.2 Fixation

A pneumothorax was applied to deeply anesthetized rats by diaphragm puncture and lungs were fixed by intratracheal application of ice-cold 2.5% glutaraldehyde in 0.15 M HEPES [13, 14]. The lungs were then dissected out, immersed in fixative for 1 day, and chopped into small cubes measuring roughly 1 mm^3^. A thoracotomy was performed on deeply anesthetized adult Wistar rats. A piece of ventricle of approximately 1 mm^3^ was excised and immediately immersed in ice-cold 2.5% glutaraldehyde in 0.15 M Na-cacodylate solution, for 1 day. Deeply anesthetized mice were perfused with 0.1 M PBS and then with 2.5% glutaraldehyde 2% paraformaldehyde in 0.1 M Na-cacodylate (all solution with pH 7.4). The cerebellum was isolated and immersed in the same solution overnight [15]. 180 μm sections were produced with a vibratome.

### 5.3 Staining

The samples were stained following one of the two protocols described below.

#### 5.3.1 Staining protocol with uranyl acetate

This protocol is known as reduced osmium, thiocarbohydrazide, osmium (ROTO) [16, 17]. Samples were rinsed 3x5 min in ice-cold 0.15 M Na-cacodylate. They were then incubated in 0.15 M Na-cacodylate solution containing 2% OsO_4_ and 1.5% potassium ferrocyanide for 1 h at 4 °C. They were rinsed 3×5 min with water, incubated for 20 min in 1% thiocarbohydrazide, rinsed again 3 × 5 min with water, and incubated in 2% OsO4 for 30 min. They were rinsed 375 min with water, incubated overnight at 4 °C in 1% uranyl acetate, rinsed 3 × 5 min with water, incubated in 1% Walton's lead aspartate [18] at 60 °C for 30 min, and rinsed 3 × 5 min with water.

#### 5.3.2 Staining protocol with lanthanides

Unless otherwise specified, the protocol based on lanthanides was the following. Samples were rinsed 3×5 min in ice-cold 0.15 M Na-cacodylate. They were then incubated in 0.15 M Na-cacodylate solution containing 2% OsO_4_ and 1.5% potassium ferrocyanide for 45 min at room temperature and for 15 min in a water bath at 50 °C. They were rinsed 3 × 5 min in water. They were incubated with 0.64 M pyrogallol for 15 minutes at room temperature and for 5 minutes in a water bath at 50 °C, and subsequently rinsed 3×5 min with water. Osmium amplification by pyrogallol instead of TCH was shown to generate stronger staining in the core of extremely large samples (whole mouse brain) [19]. Note that all the samples that we investigated were considerably thinner. The samples were incubated in 2% OsO_4_ for 22 min at room temperature and 8 min in a water bath at 50 ^×^C. After 3°5 min rinses in water, the samples were incubated overnight in a solution of 0.15 M gadolinium acetate (LFG Distribution, Lyon, France) and 0.15 M samarium acetate (LFG Distribution) pH 7.0. They were then rinsed 3×5 min with water, incubated in 1% Walton's lead aspartate [18] at 60 °C for 30 min, and rinsed 3×5 min with water.

### 5.4 Dehydration, embedding, and mounting

After staining, the samples were dehydrated in a graded ethanol series (20%, 50%, 70%, 90%, 100%, 100%) at 4°C, each step lasting 5 min. They were then infiltrated with Durcupan resin mixed with ethanol at ratios of 1:3 (v/v), 1:1, and 3:1, each step lasting 2 h. They were then infiltrated with pure Durcupan overnight. The samples were transferred to fresh Durcupan and the resin was polymerized for 3 days at 60 °C. Sample blocks were mounted on aluminum pins (Gatan, Pleasonton, CA, USA) with a conductive epoxy glue (CW2400, Circuitworks, Kennesaw, GA, USA). Care was taken to have osmicated material directly exposed at the block surface in contact with the glue in order to reduce specimen charging under the electron beam. Pyramids with a surface of approximately 500 × 500 μm^2^ were trimmed with a razor blade.

### 5.5 Electron microscopy

SBFSEM was performed with a Quanta FEG 250 microscope (FEI, Eindhoven, The Netherlands), equipped with a 3View2XP in situ ultramicrotome (Gatan). Section thickness was set between 50 and 100 nm. Images were acquired with magnifications and image sizes resulting in pixel sizes at specimen level ranging from 3 to 40 nm. Images were acquired in high vacuum mode (1–4 mPa), except where indicated otherwise. Acceleration voltage ranged from 3 to 5 kV and pixel dwell time was set between 0.5 and 16 μs.

### 5.6 Data availability

The original images used in Supplementary Video S1 will be available on the EMPIAR repository <u>(http://www.ebi.ac.uk/pdbe/emdb/empiar/)</u>. All other datasets generated during and/or analysed during the current study are available from the corresponding author on reasonable request.

## 6 Acknowledgment

This project was supported by a Swiss National Science Foundation grant (#163761 to BZ). Imaging was performed on equipment supported by the Microscopy Imaging Center (MIC), University of Bern, Switzerland.

## 7 Author contributions

A. O. designed the project, performed experiments, and analyzed data. J. L., J. R., C. R., and S. T. performed experiments. S. S and S. R contributed materials and analyzed data. B. Z designed, directed and supervised the project, analyzed data, and wrote the manuscript with feedback from all authors.

## 8 Supplementary material

Supplementary Video S1. 200-slices stack through lanthanide-stained cerebellum Purkinje layer. The sample presents no cutting damage. Figure 1D was rendered from a part of this stack. Field of view size: 10.414 × 10.913 μm^2^.

